# Hi-C-LSTM: Learning representations of chromatin contacts using a recurrent neural network identifies genomic drivers of conformation

**DOI:** 10.1101/2021.08.26.457856

**Authors:** Kevin B. Dsouza, Alexandra Maslova, Ediem Al-Jibury, Matthias Merkenschlager, Vijay K. Bhargava, Maxwell W. Libbrecht

## Abstract

Despite the availability of chromatin conformation capture experiments, understanding the relationship between regulatory elements and conformation remains a challenge. We propose Hi-C-LSTM, a method that produces low-dimensional latent representations that summarize intra-chromosomal Hi-C contacts via a recurrent long short-term memory (LSTM) neural network model. We find that these representations contain all the information needed to recreate the original Hi-C matrix with high accuracy, outperforming existing methods. These representations enable the identification of a variety of conformation-defining genomic elements, including nuclear compartments and conformation-related transcription factors. They furthermore enable in-silico perturbation experiments that measure the influence of cis-regulatory elements on conformation.

## Background

The organisation of the genome in 3D space inside the nucleus is important to its function. Chromosome conformation capture (3C) techniques, developed in the last couple of decades, have enabled researchers to quantify the strength of interactions between loci that are nearby in space. Hi-C [1] uses a combination of chromatin conformation capture and high-throughput sequencing to assay pairwise chromatin interactions genome-wide. This rich source of data promises to help elucidate genome function but its size and complexity necessitates the development computational tools.

Representation learning [2] is a machine learning technique that aims to summarize high dimensional datasets into a low-dimensional representation. It has become a valuable tool for finding compact and informative representations that disentangle explanatory factors in diverse data types. Representation learning has recently driven advances in a variety of tasks including speech recognition [3], signal processing [4], object recognition [5], natural language processing [6, 7] and domain adaptation [8]. Representation learning has recently been applied to genomic sequences [9, 10] and Hi-C data [11, 12, 13, 14].

We need representations for Hi-C data that can effectively summarize the contact map. Such a representation would encapsulate all the contacts from each locus to the others into a small number of features per position. Reducing the Hi-C map to locus-level representations in this way would allow us to study the effect of sequence elements on chromatin conformation, identify genomic drivers of 3D conformation and predict the effect of genetic variants.

Two methods for representation learning of Hi-C data have previously been developed, SNIPER [11] and SCI [12] (section Related Work). SNIPER uses a fully-connected autoencoder [15] to transform the sparse Hi-C inter-chromosomal matrix into a dense one row-wise, the bottleneck of which is assigned as the representation for the corresponding row. SCI [12] treats the Hi-C matrix as a graph and performs graph embedding [16], aiming to preserve the local and the global structures to form representations for each node.

Existing methods for Hi-C representations have two weaknesses that limit their applicability. First, SNIPER takes only inter-chromosomal contacts as input and therefore its representations cannot incorporate intra-chromosomal contact patterns such as topological domains and promoter-enhancer looping. Second, the Hi-C representations produced by both SNIPER and SCI do not account for the inherent sequential nature of the genome.

In this work, we propose a method called Hi-C-LSTM that produces low-dimensional representations of the Hi-C intra-chromosomal contacts, assigning a vector of features to each genomic position that represents that position’s contact activity with all other positions in the given chromosome. Hi-C-LSTM defines these representations using a sequential long short-term memory (LSTM) neural network model which, in contrast to existing methods like SNIPER and SCI, accounts for the sequential nature of the genome. A second methodological innovation of Hi-C-LSTM is that, instead of learning an encoder to create representations, we learn our representations directly through iterative optimization. We find that this approach provides a large improvement in information content relative to existing non-sequential methods, enables the use of intra-chromosomal interactions, and enables the model to accurately predict the effects of genomic perturbations (Results).

We demonstrate the utility of Hi-C-LSTM’s representations through several analyses. First, we show that our representations have information needed to recreate the Hi-C matrix and that this recreation is more accurate using an LSTM than alternatives. Second, we show that our representation captures cell type-specific functional activity, genomic elements and identifies genomic regions that drive conformation. Third, we show that feature attribution of Hi-C-LSTM can identify sequence elements driving 3D conformation, such as binding sites of CTCF and cohesin subunits [17, 18]. Fourth, we show that in-silico perturbation of CTCF and cohesin binding sites has the expected effects on predicted contacts, demonstrating Hi-C-LSTM’s utility for such experiments.

### Related work

Hi-C-LSTM performs two main tasks; it forms Hi-C representations, and it predicts Hi-C contacts. Learning methods have been proposed that perform either of these tasks. SNIPER [11] and SCI [12] can form representations of Hi-C. SNIPER forms Hi-C representations using a feed-forward neural network autoencoder. While SNIPER predicts high-resolution Hi-C contacts using low-resolution contacts as input, Hi-C-LSTM predicts Hi-C contacts using just the genomic positions as input. SCI forms Hi-C representations by performing graph network embedding on the Hi-C data. SCI is similar to Hi-C-LSTM in that it can be used to identify elements, however, it differs in the underlying structure it uses to represent the genome. SCI represents the genome using a graph, whereas Hi-C-LSTM treats the genome as a sequence. We compare Hi-C-LSTM with these two methods as they are most similar to what we are trying to achieve.

The first Hi-C representations were formed using Principal component analysis (PCA) based methods, introduced in Lieberman-Aiden et al. [1]. These methods cluster the Hi-C matrix into A and B compartments based on the first principal component of the intra-chromosomal contact matrix. Imakaev et al. [19] later showed that PCA based reduction is inaccurate at classifying compartments and Rao et al. [20] used a Gaussian hidden Markov model (HMM) to obtain latent features that were better at locating compartments. We treat the PCA based method developed in Lieberman-Aiden et al. [1] as a baseline.

Some methods form chromatin representations but are not directly comparable to ours. REACH-3D [21] forms internal Hi-C representations using manifold learning combined with recurrent autoencoders, however, these are three dimensional and mainly used for 3D chromatin structure inference. MATCHA [14] forms representations using hypergraph representation learning and uses them to distinguish multi-way interactions from pairwise interaction cliques. We don’t compare Hi-C-LSTM with MATCHA because MATCHA works with multi-way interaction data (SPRITE and ChIA-Drop) whereas we use pair-wise interaction data (Hi-C).

Many methods have been proposed for predicting Hi-C contacts. Some methods try to predict the chromatin contacts by using either the nucleotide sequence or chromatin accessibility and histone modifications or both [22, 23, 24, 25, 26, 27]. Akita in particular [27], is a convolutional neural network that predicts chromatin contacts from the nucleotide sequence alone, and can be used to perform in-silico predictions. In addition to these, the maximum entropy genomic annotation from biomarkers associated to structural ensembles (MEGABASE) coupled with an energy landscape model for chromatin organization called minimal chromatin model (MiChroM), generates an ensemble of 3D chromosome conformations [28]. Though these methods are similar to Hi-C-LSTM in that they predict Hi-C contacts, we don’t compare Hi-C-LSTM with them as none of them produce Hi-C representations.

## Results

### Hi-C-LSTM representations capture the information needed to create the Hi-C matrix

Hi-C-LSTM assigns a representation to each genomic position in the Hi-C contact map, such that a LSTM [29] that takes these representations as input can predict the original contact map (Fig. 2). The representation and the LSTM are jointly trained to optimize the reconstruction of the Hi-C map. This process gives us position-specific representations genome-wide (see Methods for more details).

**Figure 1:**
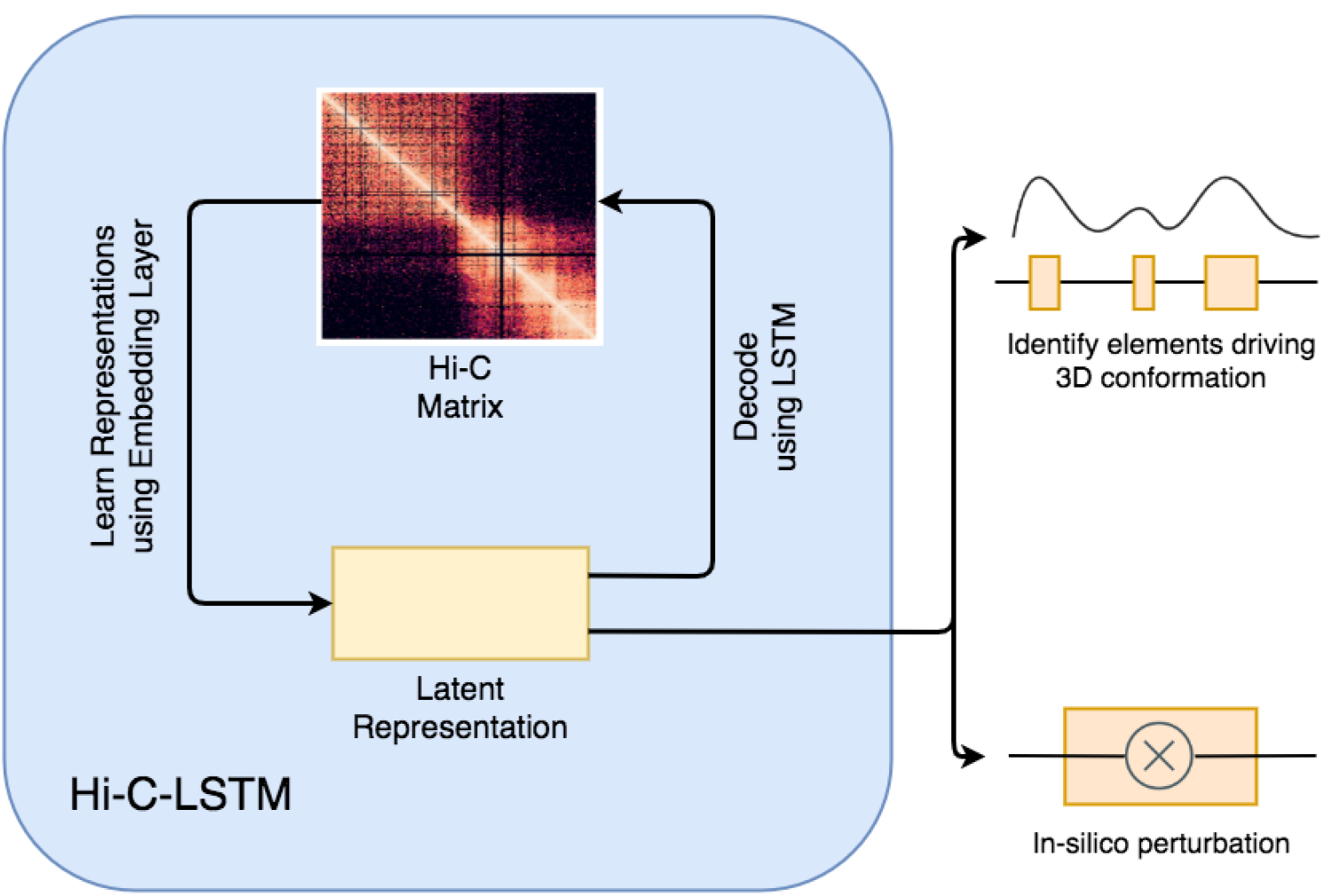
Overview of approach. Hi-C-LSTM learns a *K*-length vector representation of each genomic position that summarizes its chromatin contacts, using an LSTM embedding neural network. The representations and LSTM decoder are jointly optimized to maximize the accuracy with which the decoder can reproduce the original Hi-C matrix given just the representations. The resulting representations identify sequence elements driving 3D conformation through Integrated Gradients (IG) analysis, and they enable a researcher to perform in-silico perturbation experiments by editing the representations and observing the effect on predicted contacts.

**Figure 2:**
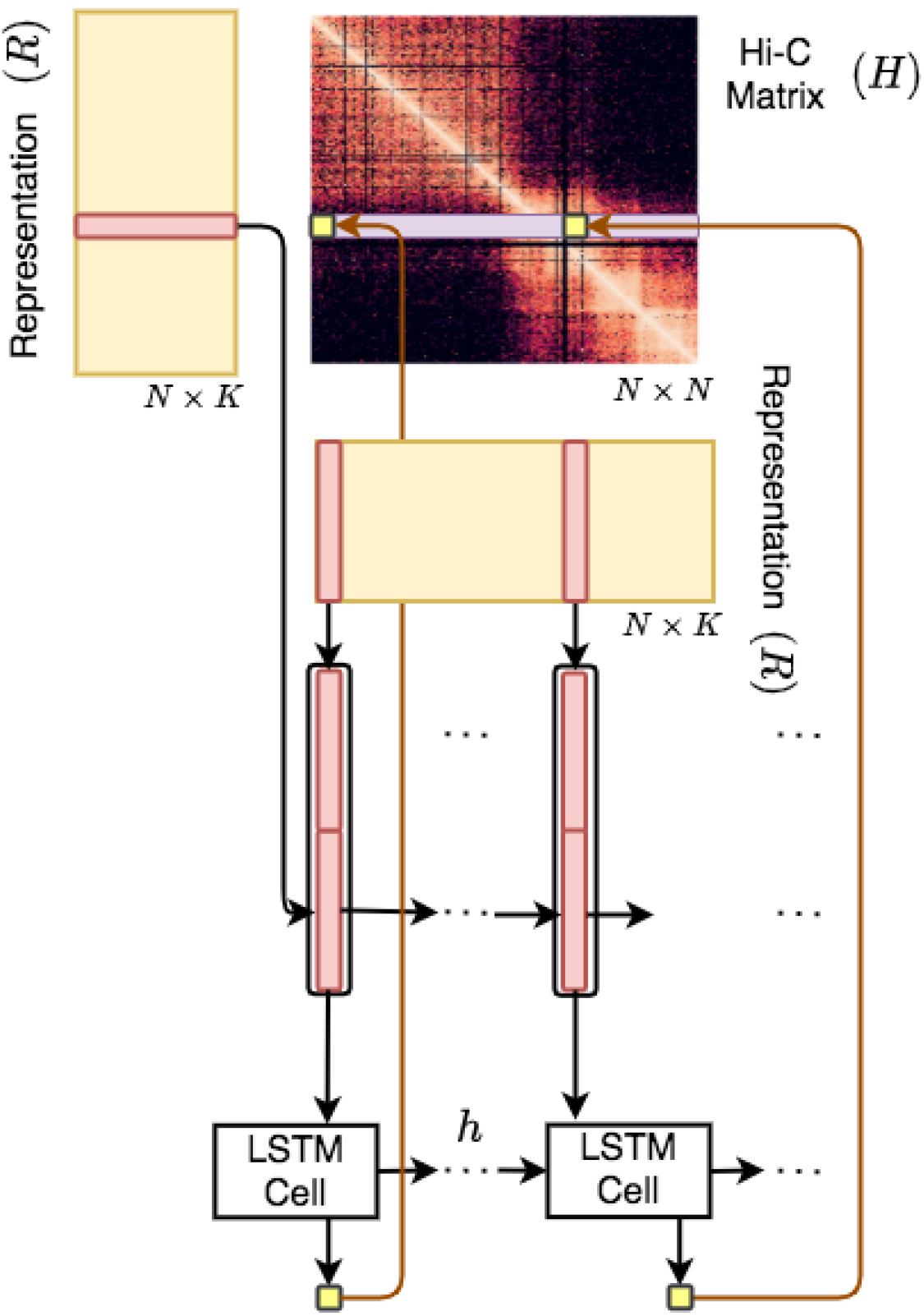
Overview of the Hi-C-LSTM model. A trained Hi-C-LSTM model consists of a *K*-length representation *R*_*i*_ for each genomic position *i* and LSTM connection weights (Methods). To predict the contact vector of a position *i* with all other positions, the LSTM iterates across the positions *j* ∈ {1 … *N*}. For each (*i, j*) pair, the LSTM takes as input the concatenated representation vector (*R*_*i*_, *R*_*j*_) and outputs the predicted Hi-C contact probability *H*_*i,j*_. The LSTM hidden state *h* is carried over from (*i, j*) to (*i, j* + 1). This process is repeated for all *N* rows of the contact map by reinitializing the LSTM states. The LSTM and the representation matrix are jointly trained to minimize the reconstruction error.

We find that the Hi-C-LSTM achieves higher accuracy when constructing the Hi-C matrix compared to existing methods (Fig. 3A). The inferred Hi-C map matches the original Hi-C map (Fig. 3C) closely, and differs from it by about 0.25 R-squared points on average. We adapt SNIPER to our task by replacing the feed-forward decoder that converts low-resolution Hi-C to high-resolution Hi-C with a decoder that reproduces the original input Hi-C. We call this SNIPER-FC. Hi-C-LSTM outperforms SNIPER (SNIPER-FC) convincingly, by 10% higher R-squared on average (Fig. 3A). Hi-C-LSTM also outperforms SCI (SCI-LSTM) by 12% higher R-squared on average (Fig. 3A).

**Figure 3:**
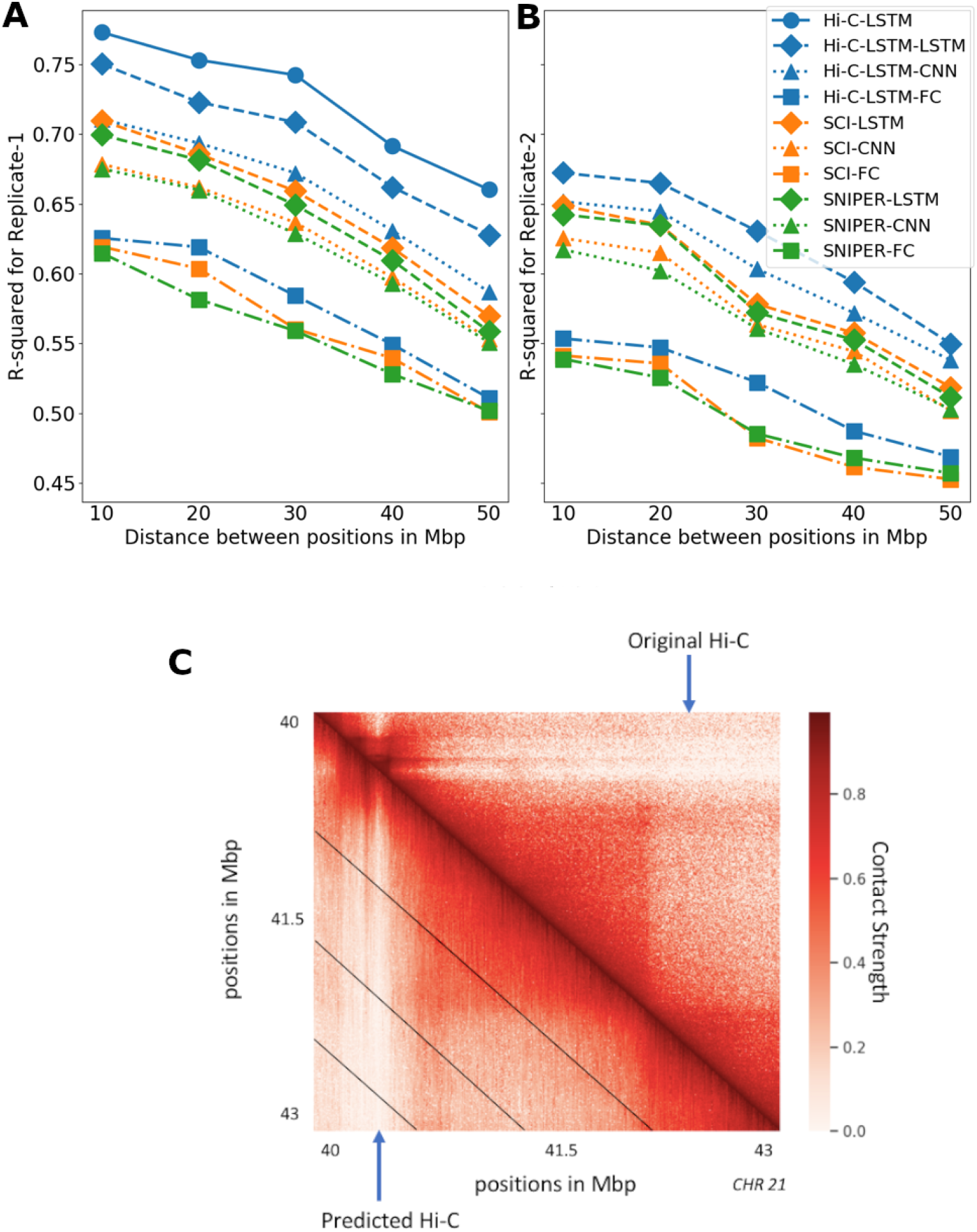
Accuracy with which representations reproduce the original Hi-C matrix. **A**,**B**) The Hi-C R-squared computed using the combinations of representations from different methods and selected decoders for replicate 1 and 2. The horizontal axis represents the distance between positions in Mbp. The vertical axis shows the average R-squared for the predicted Hi-C data. The R-squared was computed on a test set of chromosomes using selected decoders with the representations trained all chromosomes as input. The legend shows the different combinations of methods and decoders, read as *[representation]-[decoder]*. **C**) A selected portion of the original Hi-C map (upper-triangle) and the predicted Hi-C map (lower-triangle). The portion is selected from chromosome 21, between 40 Mbp to 43 Mbp. Diagonal black lines denote Hi-C-LSTM’s frame boundaries (Methods).

Two hypotheses could explain Hi-C-LSTM’s improved reconstructions: (1) that Hi-C-LSTM’s representation captures more information, or (2) that an LSTM is a more powerful decoder. We found that both are true. To distinguish these hypotheses, we split each model respectively into two components—its representation and decoder—and evaluated each possible pair of components. We train the representations (Hi-C-LSTM, SCI, SNIPER) on all chromosomes and couple them with selected decoders (LSTM, CNN, FC). Using the representations as input, we retrain these decoders with a small subset of the chromosomes and test on the rest. (see Methods for more details). We compute the average R-squared value for creating the Hi-C contact matrix using each combination of selected representations and decoders

We found that the choice of decoder has the largest influence on reconstruction performance. Using a LSTM decoder performs best, even when using representations derived from SNIPER or SCI (improvement of 0.14 and 0.12 R-squared points on average over fully-connected decoders respectively, Fig. 3A). Furthermore, we found that Hi-C-LSTM’s representations are most informative, even when using decoder architectures derived from SNIPER or SCI (Fig. 3A).

Though the Hi-C-LSTM representations capture important information from a particular sample, we wanted to verify whether they capture real biological processes or irreplicable experimental noise. To check the effectiveness of Hi-C-LSTM representations in creating the Hi-C contact map of a biological replicate, we train the representations on one replicate (replicate 1), repeat the decoder training process on replicate 2 (see Methods for more details), and compute the average R-squared value for creating the Hi-C contact map of replicate 2 (Fig. 3B). The average R-squared reduces slightly for inference of replicate 2 due to experimental variability; however, the performance trend of the representation-decoder combinations is largely preserved (Fig. 3B). These results show that Hi-C-LSTM’s improved performance is not merely driven by memorizing irreplicable noise.

### Hi-C-LSTM representations locate functional activity, genomic elements, and regions that drive 3D conformation

Considering that a good representation of Hi-C should contain information about the regulatory state of genomic loci, we evaluated our model by checking whether these genomic phenomena and regions are predictable from only the representation. Specifically, we test whether the position specific representations learned via the Hi-C contact-generation process are useful for genomic tasks that the model was not trained on, such as classifying genomic phenomena like gene expression [30] and replication timing [31, 32, 33, 34], locating nuclear elements like enhancers, transcription start sites (TSSs) [35] and nuclear regions that are associated with 3D conformation like promoter-enhancer interactions (PEIs) [36, 37, 38], frequently interacting regions (FIREs) [39, 40], domains, loops and subcompartments [20]. We used a boosted decision tree (XGBoost) model [41] to predict binary genomic features from representations. (See Methods for more details regarding comparison methods, baselines and classifier).

We find that the models built using the intra-chromosomal representations achieve higher predictive accuracy overall relative to ones trained on inter-chromosomal representations when predicting gene expression, enhancers and TSSs (Fig. 4A). This trend is likely due to the relatively close range of the elements involved in prediction. In contrast, SNIPER is slightly better at predicting replication timing when compared to the rest of the intra-chromosomal models except Hi-C LSTM (SNIPER-INTER, Fig. 4A). While all methods achieve low absolute accuracy at predicting promoter-enhancer interactions, Hi-C-LSTM performs best (0.5 mAP on average, 0.1 mAP higher on average than SCI) (Fig. 4A, B). Both methods perform comparably in predicting the other interacting genomic regions like FIREs, domains, loops, and subcompartments (Fig. 4A). SNIPER-INTRA as well as SNIPER-INTER don’t perform as well as Hi-C-LSTM and SCI on these tasks.

**Figure 4:**
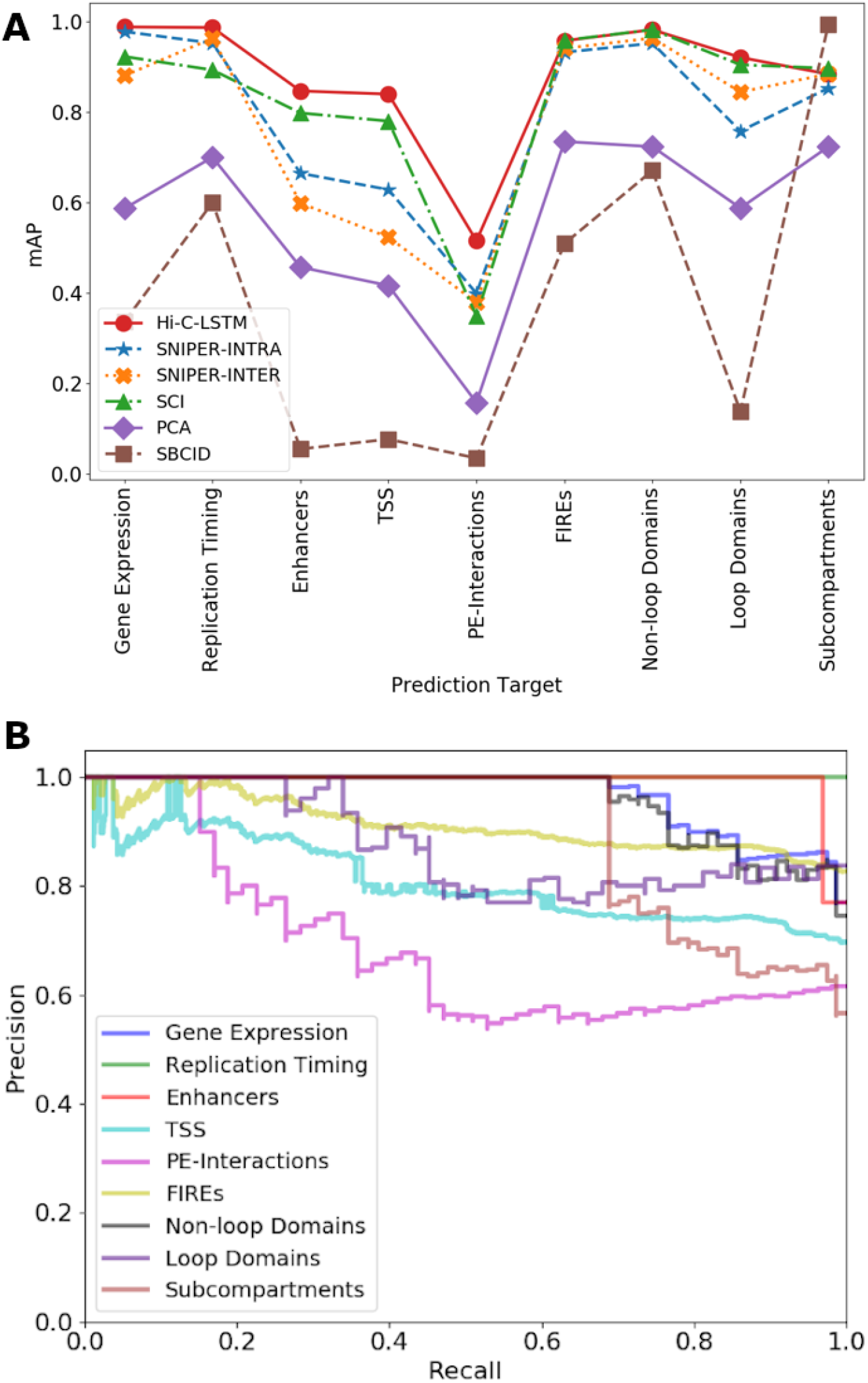
Important genomic phenomena and chromatin regions are classified using the Hi-C-LSTM representations as input. **A**) mAP for gene expression, replication timing, enhancers, transcription start sites (TSSs), promoter-enhancer interactions(PEIs), frequently interacting regions (FIREs), loop and non-loop domains, and subcompartments. The y-axis shows the mAP, the x-ticks refer to the prediction targets, and the legend shows the different methods compared with. **B**) The Precision-Recall curves of Hi-C-LSTM for the various prediction targets. The y-axis shows the Precision, the x-axis shows the Recall, and the legend shows the prediction targets.

The only task on which other methods outperform Hi-C-LSTM is at predicting subcompartments. Subcompartments were originally defined based on inter-chromosomal interactions, so representations based on such interactions outperform those based on intra-chromosomal interactions such as Hi-C-LSTM. Also subcompartment-ID (SBCID) (Methods) achieves perfect mAP by virtue of its design (Fig. 4A). Among the rest of the methods, we find that methods which were designed to predict subcompartments such as SCI and SNIPER-INTER, perform better than the others (Fig. 4A). Hi-C-LSTM does perform marginally better than SNIPER-INTRA. Overall, although Hi-C LSTM performs better than other models on most of the tasks, the performance of SCI and SNIPER are comparable to Hi-C-LSTM and all three models perform significantly better than the baselines on average (Fig. 4A).

### Feature attribution reveals association with genomic elements driving 3D conformation

Given that our representations capture elements driving 3D conformation, we should be able to identify those elements using our representations. To validate the ability of our representations to locate genomic regions that drive chromatin conformation, we identified which genomic positions have the largest impact on Hi-C contacts, using the technique of feature attribution. Feature attribution is a technique that allows us to attribute the prediction of neural networks to their input features. In this case, it identifies which genomic positions influence which Hi-C contacts. We ran feature attribution analysis on the Hi-C-LSTM and aggregated the feature importance scores across all the dimensions of the input representation to get a single score for each genomic position (see Methods for more details). We expected to see higher feature attribution for the elements, regions, and domains that are crucial for chromatin conformation.

We found that the CTCF and cohesin binding sites as given by ChIP-seq have a large influence on contacts given their high feature importance score. The genome folds to form “loop domains”, which are found to be a result of tethering between two loci bound by CTCF and cohesin subunits RAD21 and SMC3 [18]. Among the many models of genome folding, a CTCF protein- and cohesin ring-associated complex that extrudes chromatin fibers is most promising. This extrusion model explains why loops don’t overlap [17]. We found that CTCF sites show 10% higher mean importance score than RAD21 and SMC3 sites and all three sites have a spread that is predominantly positive (Fig. 5C). The high feature importance scores observed at CTCF and cohesin binding sites validates the crucial role they play in loop formation [17, 18].

**Figure 5:**
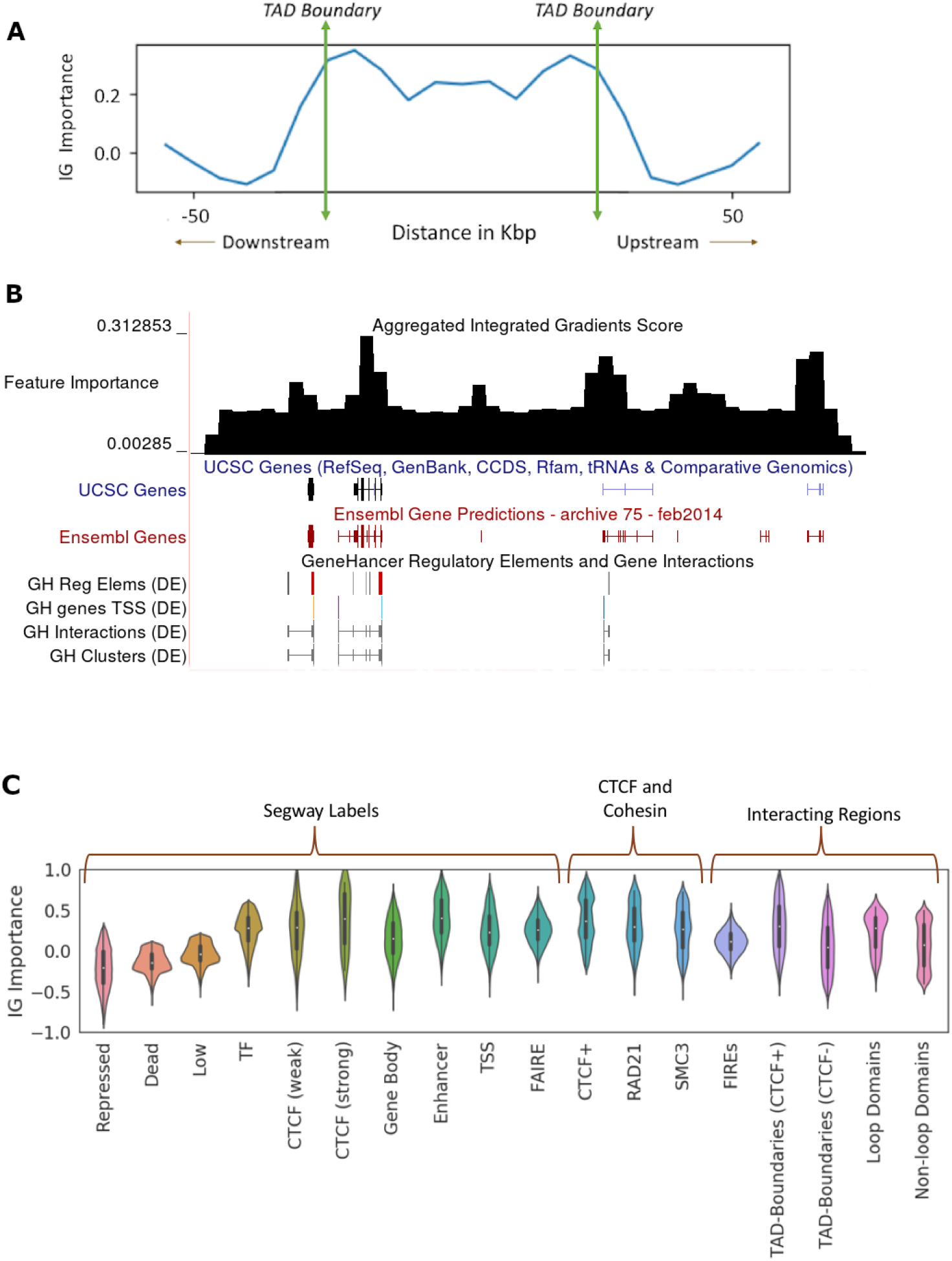
Hi-C-LSTM representations identify genomic elements involved in conformation through Integrated Gradients (IG) feature importance analysis. **A**) The IG feature importance averaged across different TADs of varying sizes. The vertical axis indicates the average IG importance at each position and the horizontal axis refers to relative distance between positions in Kbp, upstream/downstream of the TADs. **B**) The IG feature importance for a selected genomic locus (chr21 28-29.2Mbp) along with genes, regulatory elements and Hi-C. We see that the feature importance scores peak at known regulatory elements. **C**) Violin plots of aggregated feature attribution scores for selected elements. The x-axis shows the labels/elements and the y-axis displays the log plus z normalized feature importance scores from Integrated Gradients.

The importance of CTCF is further validated by the aggregated feature importance (Fig. 5C), showing a markedly positive score near CTCF binding sites given by Segway [42], particularly the strong ones (mean importance score of 0.45). Moreover, we see that the model places high importance on regulatory elements, particularly enhancers (mean importance score of 0.4) (Fig. 5C). The active domain types have a higher mean score and a spread that largely occupies the positive portion of the feature importance plot when compared to the inactive regions (Fig. 5C). This suggests that active regions may play a dominant role in nuclear organization, where the movement of repressed regions to the periphery is a side-effect.

Aggregated feature importance also demonstrates the largely positive feature attribution of genomic regions that are an integral part of 3D conformation like FIREs, topologically associating domain (TAD) boundaries with and without CTCF sites, loop and non-loop domains (Fig. 5C). TAD boundaries enriched with CTCF show a 20% higher mean importance score compared to TAD boundaries not associated with CTCF, pointing to the importance of CTCF sites at domain boundaries in conformation (Fig. 5C). Moreover, loop domains show a 20% higher mean importance score compared to non-loop domains, which is expected because of the increased contact strength on average and the presence of CTCF sites (Fig. 5C).

The variation of the aggregated feature importance across interesting genomic regions helps us distinguish boundaries of domains and genomic regulatory elements (Fig. 5). We observe the variation of the feature importance signal across TADs and a selected portion of chromosome 21 (28 Mbp to 29.2 Mbp) [43] to check if we can isolate the boundaries of domains, genes and other regulatory elements. To deal with TADs of varying sizes, we partition the interior of all TADs into 10 equi-spaced bins and average the feature importance signal within these bins. We plot this signal along with the signal outside the TAD boundary 50Kbp upstream and downstream, averaged across all TADs (Fig. 5A). The feature importance has largely similar values in the interior of the TAD, noticeably peaks at the TAD boundaries, and slopes downward in the immediate exterior vicinity of the TAD (Fig. 5A). This trend validates the importance of TADs and TAD boundaries in chromatin conformation, which we saw in (Fig. 5C). We also consider a candidate region in chromosome 21 that is referred to in [43] to observe the variation of feature importance across active genomic elements (Fig. 5B). For this selected region in chromosome 21, as we don’t have to deal with domains of varying sizes, we just average the feature importance signal within a specified number of bins and plot this in the UCSC Genome Browser [44] along with genes and regulatory elements. The feature importance peaks around genes, regulatory elements and domain boundaries (Fig. 5B), showing that they play a more important role in conformation than other functional elements.

### Hi-C-LSTM enables in-silico knockout experiments

As Hi-C-LSTM models the dependence of sequence on 3D conformation, it enables us to perform in-silico deletion, insertion and reversal of certain genomic loci and observe changes in the resulting Hi-C contact map. In-silico knockout experiments have gained prominence lately, mainly in intercepting signal flows in signaling pathways [45] and drug discovery [46, 47, 48]. A Hi-C in-silico manipulation tool is of great value it enables researchers to identify the importance and influence of any genomic locus of interest to 3D chromatin conformation.

Hi-C-LSTM enables a researcher to perform two types of experiments. First, one can simulate the knockout of a locus by deleting a portion of the representation or replacing it with a null representation. As a null, we use the average local features within 0.2 Mbp. Second, one can simulate the replacement or translocation of an element by replacing or removing the corresponding representation (see Methods).

Previous work showed that inserting even a single base pair near the loop anchors can make many loops and domains vanish, altering chromatin conformation at the megabase scale [17]. Given the crucial role played by CTCF and cohesin subunits in conformation at loop anchors (See Downstream Classification, Feature Attribution), we hypothesized that knocking out CTCF and cohesin subunit binding sites will change the Hi-C contact map noticeably. The average difference in predicted contact strength between no knockout and knockout at the site under consideration as a function of genomic distance is observed (Fig. 6C). After CTCF and cohesin knockouts, the average contact strength reduces by *>*15% when compared to the no knockout case (Fig. 6C). CTCF knockout is seen to affect insulation at about 100 Kbp and reflect possible loss of loops at 200 Kbp (Fig. 6C). The knockout of cohesin subunits SMC3 and RAD21 binding sites is observed to be independent of CTCF knockout with 5% higher average inferred strength over distance, hinting at their relative importance (Fig. 6C).

**Figure 6:**
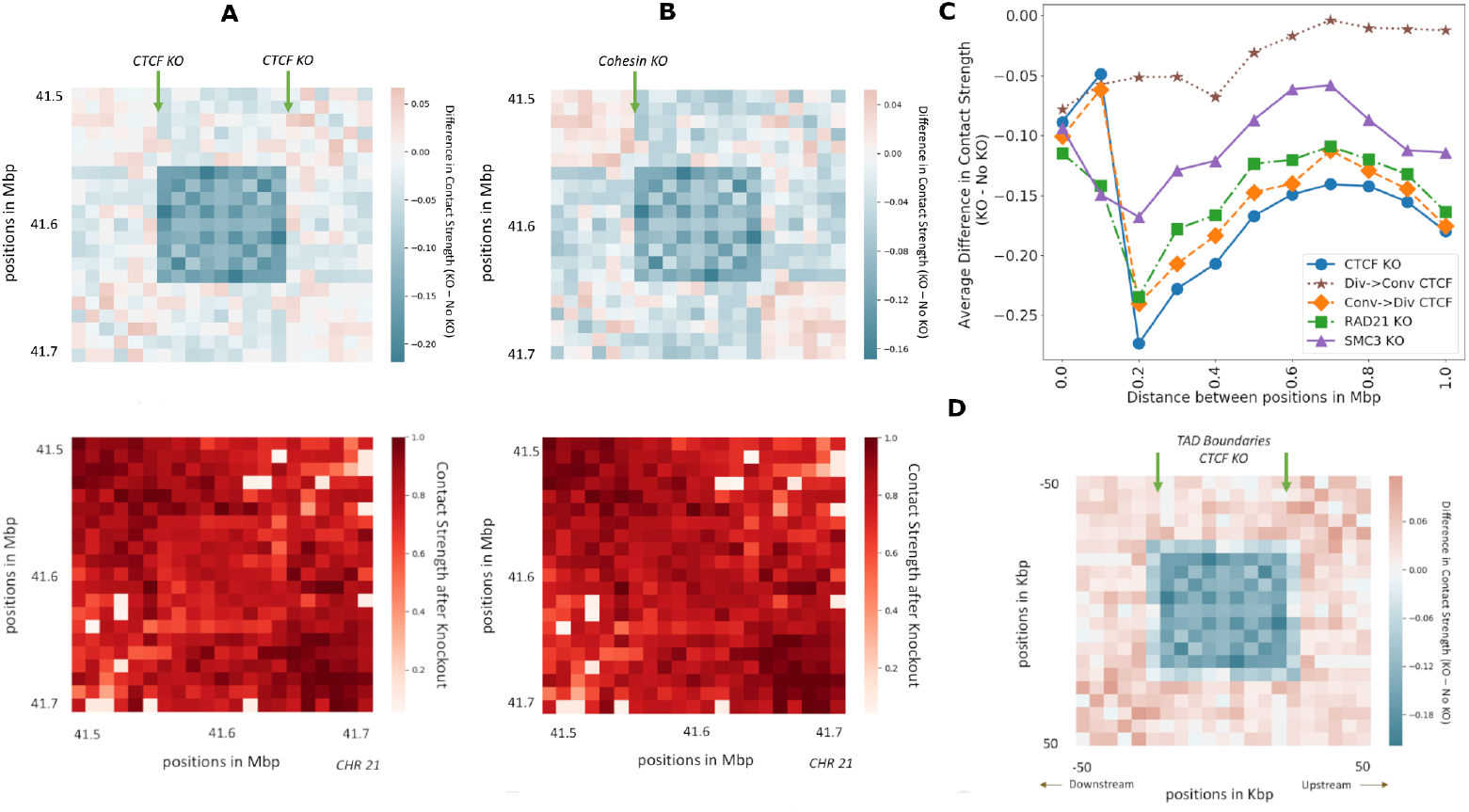
In-silico deletion and orientation replacement of CTCF and cohesin subunits is performed and changes in the resulting Hi-C contact matrix is observed. **A**) The difference in predicted Hi-C contact strength between CTCF knockout and no knockout, performed on chromosome 21, between 41.5 Mbp to 41.7Mbp. The bottom figure shows just the predicted Hi-C after knockout. **B**) Same as *A*, but for Cohesin knockout. **C**) Average difference in contact strength of the inferred Hi-C matrix between knockout and no knockout (y-axis) for varying distance between positions in Mbp (x-axis). The knockout experiments include CTCF and cohesin knockout and convergent/divergent CTCF replacements (legend). **D**) The genome-wide average difference in predicted Hi-C contact strength between CTCF knockout at TAD boundaries and no knockout.

The CTCF sites at loop anchors occur mainly in a convergent orientation, with the forward and reverse motifs together, suggesting that this formation maybe required for loop formation [20, 49, 50, 51, 52, 53, 54]. To check how important the orientation of CTCF motifs is to conformation, we conducted CTCF orientation replacement experiments at loop boundaries. The average difference in predicted contact strength between no replacement and replacement at the site under consideration as a function of genomic distance is observed (Fig. 6C). The replacement of convergent with the divergent orientation around loops is seen to behave similar to the case of CTCF knockout thereby validating observations made in [55] (Fig. 6C). On the other hand, replacement of divergent with the convergent orientation is seen to preserve loops at 200 Kbp and behave similar to the control, although with reduced inferred contact strength (5% on average) (Fig. 6C).

The difference in inferred Hi-C between the CTCF (Fig. 6A) and cohesin (Fig. 6B) knockout and the no knockout for a selected portion of chromosome 21 (41.5 Mbp to 41.7Mbp), shows the importance of CTCF and cohesin sites in conformation. The CTCF knockout at both the edges of the loop results in decrease in contact strength (0.18 lower on average) within the loop (Fig. 6A). Cohesin knockout at the start of the loop also results in decrease in contact strength within the loop (0.12 lower on average), but not as strongly as the CTCF knockout (Fig. 6B). Around the loop, CTCF and cohesin knockout results in patches of decreased (0.05 lower on average) as well as increased contacts (0.05 higher on average) (Fig. 6A, B). The predicted Hi-C after CTCF and cohesin knockout (Fig. 6A, B:Bottom) validates the fading of loops. The average difference in inferred Hi-C between the CTCF knockout at TAD boundaries and the no knockout (Fig. 6D) shows similar trends, with decreased contacts (0.2 lower on average) within the TAD and increased contacts (0.08 higher on average) outside the TAD. The symmetry of the Hi-C matrix is largely preserved after the knockouts, validating the capability of Hi-C-LSTM to perform knockout experiments.

### Hi-C-LSTM accurately predicts effects of a 2.1 Mbp duplication at the SOX9 locus

To further validate Hi-C-LSTM as a tool for in-silico genome alterations, we simulated a structural variant at the SOX9 locus that was previously assayed by Melo et al. [56]. This variant was observed in an individual with Cook’s syndrome and comprises the tandem duplication of a 2.1 Mbp region on chromosome 17 that includes regulatory elements of SOX9 (chr17:67,958,880–70,085,143; GRCh37/hg19, Fig. 7A). To simulate a Hi-C experiment on a genome with this variant, we made a new Hi-C-LSTM representation matrix that includes a tandem copy of the representation at the locus in question and passed this representation matrix through the original Hi-C-LSTM decoder to produce a simulated Hi-C matrix on a post-duplication genome (Fig. 7B). Because Hi-C reads cannot be disambiguated between the two duplicated loci, we simulated mapping reads to the original hg19 reference by summing reads originating from the two copies (see Methods). We evaluated Hi-C-LSTM’s predictions according to the agreement between this predicted matrix and a Hi-C experiment performed by Melo et al. [56] (Fig. 7C).

**Figure 7:**
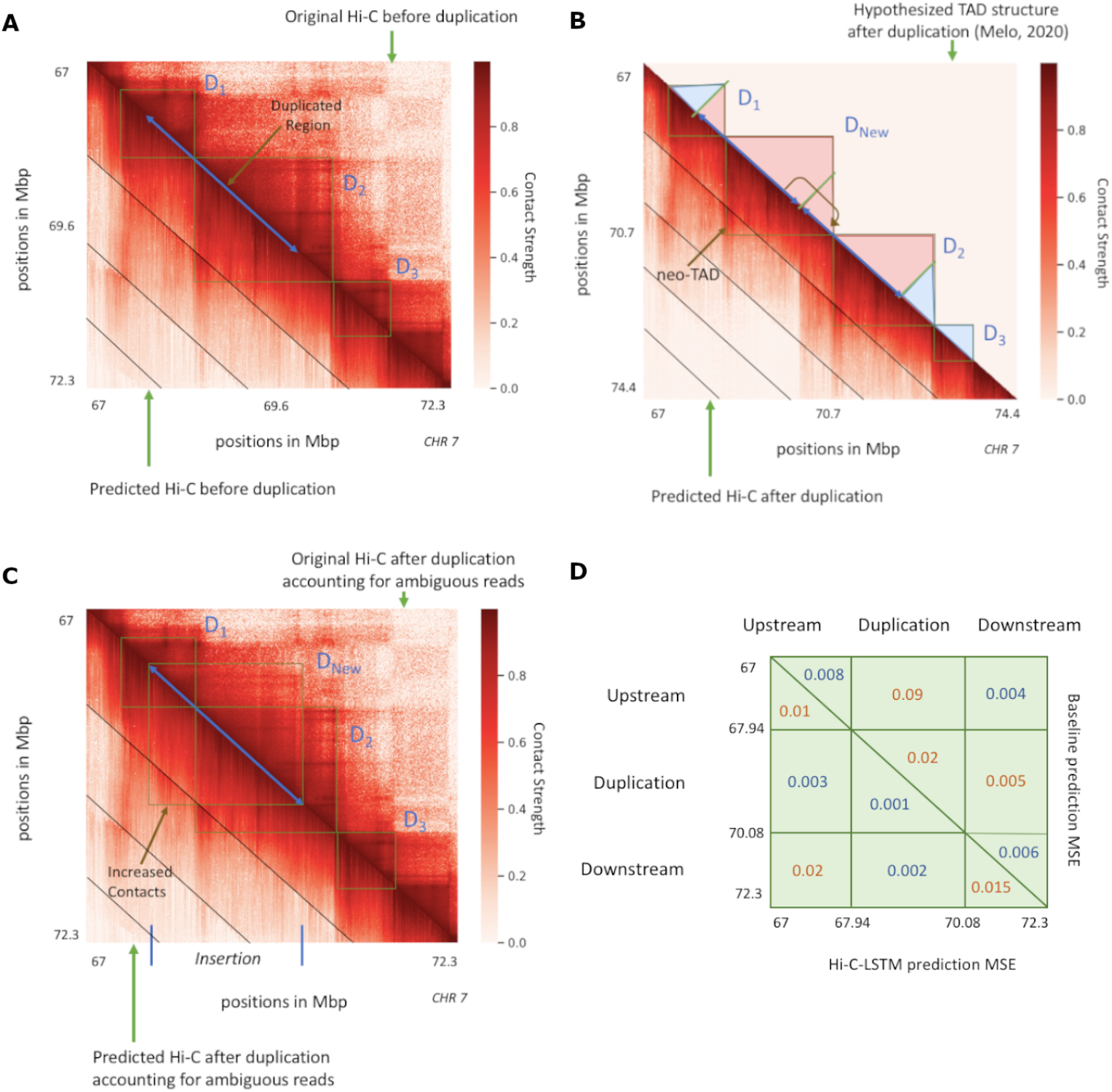
In-silico duplication of a 2.1 Mbp region on Chromosome 7 [56]. In all subplots, upper and lower triangles denote observed and predicted Hi-C contact probabilities respectively, and diagonal black lines denote Hi-C-LSTM frame boundaries. **A**) Original and predicted Hi-C before duplication. *D*_1_, *D*_2_ and *D*_3_ indicate the three pre-duplication topological domains. **B**) Predicted Hi-C after duplication on a simulated reference genome that includes both copies. Lower triangle indicates Hi-C-LSTM predicted contacts. The true Hi-C contact matrix on this reference genome is not observable because the read mapper cannot disambiguate between the two copies. The upper triangle depicts the post-duplication topological domain structure hypothesized by Melo et al, which includes a novel topological domain *D*_New_. **C**) Original and predicted Hi-C on the original pre-duplication reference genome. Upper triangle shows observed post-duplication Hi-C data assayed by Melo et al. Lower triangle shows Hi-C-LSTM predictions, mapped to the pre-duplication reference by summing the contacts for the two copies (Results). **D**) Average mean-squared error in predicting the observed data by (lower triangle) Hi-C-LSTM, and (upper triangle) a simple baseline (Results) at the upstream, duplicated, and downstream regions.

We found that Hi-C-LSTM accurately predicted the effect of the duplication. The domains that existed pre-duplication (*D*_1_, *D*_2_, *D*_3_, Fig. 7A) are correctly captured post-duplication. In addition, a new chromatin domain (*D*_New_) that was introduced by the duplication is correctly predicted by Hi-C-LSTM (Fig. 7B). To quantitatively evaluate our predictions, we compared them to a baseline that predicts the original pre-duplication Hi-C for the interactions between the upstream, downstream and duplicated regions, and the genomic average for the interactions of the duplicated region with itself (see Methods). We found that Hi-C-LSTM’s predictions significantly outperform this baseline overall (Fig. 7D). Note the baseline is a slightly better predictor of contacts between the upstream and downstream regions.

Hi-C-LSTM’s predictions have the advantage that they describe contacts on the true post-duplication genome, in contrast to the reference genome used to map reads (Fig. 7C). Hi-C-LSTM’s contacts recapitulate the post-duplication topological domain structure hypothesized by Melo et al. These duplication experiments further validate the ability of Hi-C-LSTM to perform in-silico mutagenesis.

## Discussion

In this work we have proposed a deep LSTM model that uses intra-chromosomal contacts to form position-specific representations of chromatin conformation. These representations are able to capture a variety of genomic phenomena and elements and at the same time distinguish genomic regions, transcription factors and domains that are known to play an important role in chromatin conformation. They also elucidate the interplay between genome structure and function. The classification and feature attribution results validate the ability of the representations to locate vital regions such as CTCF and cohesin binding sites.

The primary contribution of this work is the application of a deep LSTM to the problem of forming representations for intra-chromosomal interactions. The Hi-C-LSTM not only outperforms the existing models like SCI and SNIPER that form representations in predicting genomic phenomena but also locates elements driving 3D conformation as revealed by feature importance analysis. In addition to these, the Hi-C-LSTM has few distinct advantages over its counterparts. One, it can be used as a contact generation model. It’s observed that the Hi-C-LSTM representations are more informative in this regard and that sequential models like the LSTM perform much better at contact generation. Two, a low-dimensional Hi-C-LSTM representation is powerful enough to reasonably recreate the Hi-C matrix (see Ablation). Three, the Hi-C-LSTM framework allows us to conduct in-silico experiments like insertion, deletion and reversal of elements driving 3D conformation and observe changes in contact generation. This would be extremely useful in fully understanding the role of CTCF and cohesin binding sites and other transcription factors in chromatin conformation.

An important limitation of Hi-C-LSTM’s *in silico* experiment is that they can simulate only *cis* effects. Variation in chromatin structure can be caused either by *cis* or *trans* effects. *Cis* effects are caused by genetic variants on the same DNA molecule, whereas *trans* effects arise from diffusible elements like transcription factors. Hi-C-LSTM can model only *cis* effects because *trans*-acting cellular machinery is captured within the Hi-C-LSTM decoder, which cannot be easily modified. An example of a cis-effect is the duplication at the SOX9 locus, in which case we showed Hi-C-LSTM correctly models the resulting neo-TAD (see Duplication) [56]. Hi-C-LSTM cannot model *trans* effects such as recent investigation of the removal of RAD21 [18] and CTCF [57, 58].

The good performance of Hi-C-LSTM suggests several avenues for future work. First, extending the mode to incorporate data from multiple cell types and the resulting representations may yield insights into differences in chromatin organization across development. Second, the success of a LSTM model suggests trying other recurrent neural network models such as Transformers [59]. Third, a modified version of Hi-C-LSTM may be able to infer a 3D structure of chromatin. The Hi-C representations that we form currently are embedded on a lower-dimensional manifold that does not have any direct physical significance. However, a Hi-C-LSTM-like model trained to produce three-dimensional representations may be able to reproduce the true nuclear positions of chromatin.

## Conclusions

Hi-C-LSTM representations capture genomic regions that play a vital role in chromatin conformation. The utility of these representations include but is not limited to: supervised classification to find association with genomic phenomena, unsupervised element discovery using feature importance and in-silico knockout to elucidate the role of sequence in conformation.

## Methods

The code and data repository for this project, including training, evaluation, data handling, and generated data can be found in our GitHub repository [60].

### Data sets

The Hi-C data for the GM12878 B-Lymphocyte cell line was acquired using the GEO accession number GSE63525 [20, 61]. We generated a intrachromosomal Hi-C data set on the hg19 human reference genome assembly [62] at 10Kb resolution with KR (Balanced) normalization [63] using juicer tools [64] with the command `java -jar juicer_tools.jar dump observed KR data/chr.hic chr chr BP 10000 chr.txt`, where `chr` refers to the chromosome being extracted.

Following SCI [12], to mitigate the extreme range of magnitudes present in Hi-C read counts, we transformed Hi-C values into contact probabilities between 0 and 1. We calculated contact probabilities according to the exponential transformation (Eq. 1)

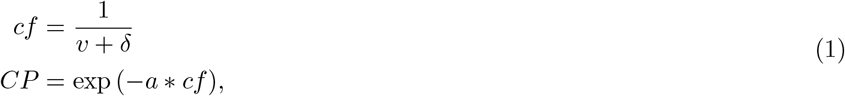

where *v* is the raw input contact strength, *δ* is a very small positive real number (we set *δ* to be 10^−10^), *cf* is the coefficient obtained, *a* is the coefficient multiplier, and *CP* is the resulting contact probability. We chose *a* = 8 because it appeared to provide a good separation of low and high contact values.

RNA-seq data for 57 cell types was obtained from the Roadmap Consortium [65].

For the classification task, each gene was considered to be active if its log mean expression value across the gene was greater than 0.5 [66, 67].

We defined promoter-enhancer interactions as the ones that were used to train TargetFinder [68, 69].

Frequently interacting region (FIRE) scores at 40Kbp resolution were downloaded from the additional material of [39] and were converted to binary indicators using 0.5 as a threshold following [70].

The replication timing data given by Repli-Seq [71] was downloaded from Replication Domain [72] at 40Kbp resolution.

Locations of known enhancers and transcription start sites (TSSs) were obtained from FANTOM [73] and ENCODE [74] respectively.

Domain, loop and subcompartment annotations were obtained from the results of Rao et al. [20] using the GEO accession number GSE63525 [61].

Segway and Segway-GBR labels were obtained from Hoffmanlab [75] and Noblelab [76] respectively.

CTCF, RAD21 and SMC3 peak calls were downloaded from ENCODE [77]. The CTCF orientations were obtained by using the CTCF motif from the MEME suite [78] (version 5.3.3) and running FIMO [79] to get the motif instances using the command `fimo -oc output_directory motif_file.meme sequence_file.fna`. We use all default options while running fimo including the p-value threshold (`--thresh`) of 10^−4^. We ran FIMO after obtaining the human genome sequence file under mammals and the hg19 genome assembly.

Topologically-associating domain (TAD) annotations were downloaded from TADKB [80].

### LSTM

Long short-term memory (LSTM) networks were proposed as a solution to the vanishing gradient problem [81] in recurrent neural networks (RNNs) [82]. They are known to be a good candidate for modelling sequential data and have been widely used for sequential tasks [83, 84, 85]. An LSTM is made up of a memory state (*h*_*t*_), a cell state (*c*_*t*_), and three gates that control the flow of data: input (*i*_*t*_), forget (*f*_*t*_) and output (*o*_*t*_) gates. The input and the forget gates together regulate the effect of a new input on the cell state. The output gate determines the contribution of the cell state on the output of the LSTM.

Let matrices *W* and *U* be the weights of the input and recurrent connections, and *b* refer to the biases. There are four sets of weight matrices and biases in the LSTM. These include one for each of the three gates—forget gate (*W*_*f*_, *U*_*f*_, *b*_*f*_), input gate (*W*_*i*_, *U*_*i*_, *b*_*i*_) and output gate (*W*_*o*_, *U*_*o*_, *b*_*o*_)—and one to form the cell state (*W*_*c*_, *U*_*c*_, *b*_*c*_). The current cell state (*c*_*t*_) is formed by the modulation of the previous cell state (*c*_*t−*1_) by the forget gate (*f*_*t*_) and combining it with the modulation of the current input (*x*_*t*_) and the previous memory state (*h*_*t−*1_) by the input gate (*i*_*t*_). Finally, the current memory state (*h*_*t*_) is formed by the modulation of the current cell state (*c*_*t*_) by the output gate (*o*_*t*_).

An LSTM’s output is determined by the following series of operations [29].

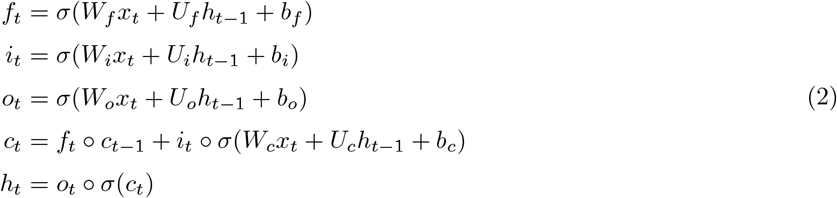

where ○ is the Hadamard product and *σ* refers to the sigmoid activation function.

### Hi-C-LSTM

Hi-C-LSTM creates a representation given a pair of genomic positions in the Hi-C contact matrix using an embedding neural network layer and predicts the contact strength at that particular pair via a deep LSTM [29] that takes these representations as input (Fig. 2). Hi-C-LSTM takes as input a *N* × *N* intra-chromosomal Hi-C contact matrix (ℍ^*N*×*N*^), for each chromosome, where *N* is the chromosome length.

A trained Hi-C-LSTM model consists of LSTM parameters (section LSTM) and a representation matrix *R* ∈ ℝ^*N*×*M*^, where *M* is the representation size. At each genomic position, (*i, j*) pair is given as input to an embedding layer, which indexes the row and column representations *R*_*i*_, *R*_*j*_ ∈ ℝ^𝕄^ and feeds these two vectors as input to the LSTM. The output of the LSTM is the predicted Hi-C contact probability *Ĥ*_*i,j*_ for the given (*i, j*) pair.

The hidden states of the LSTM are carried over from preceding columns thereby maintaining a memory for the row. For the sake of memory usage, the hidden states are reinitialized after every each frame of 1.5 Mbp or 150 resolution bins (see Design Choices). This process is repeated for each row of the Hi-C matrix (Eq. 3).

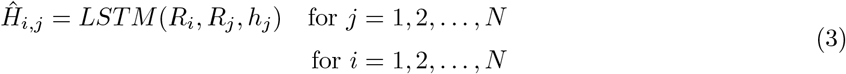

where *h*_*j*_ is the same as *h*_*j−*1_ within the frame and is reinitialized at the beginning of each new frame.

The LSTM and the embedding neural network layer are jointly trained using the mean squared error (MSE) loss function which facilitates the faithful construction of the Hi-C intra-chromosomal matrix (Eq. 4).

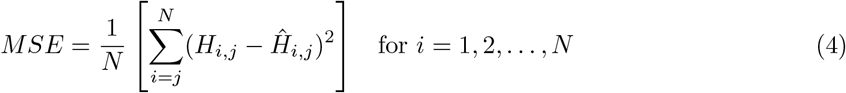

At the end of all the training iterations, the output of the embedding neural network layer at each row *i* (*R*_*i*_) is treated as the representation for that row. The Hi-C-LSTM framework infers the Hi-C contact matrix from pairs of position IDs and therefore is a transformation from linear sequential space to the Hi-C space. The linear position IDs are a convenient and useful modeling assumption which builds a framework that doesn’t make any other transfer function assumptions.

### Modeling choices and training

The LSTM model required us to make a few design choices. As layer normalization can significantly reduce the training time and is effective at stabilizing the hidden state dynamics in LSTMs, we used a unidirectional layer norm LSTM [86] with one hidden layer. We found that variants such as the bidirectional LSTM [87] and LSTM with multiple layers provided a marginal increase in test performance (Additional Files: Fig. 1). The variants were also prone to overfitting. Therefore, we chose the single-layer unidirectional model over these variants accounting for computational efficiency and good generalization. Gradient clipping [81] and the *softsign* activation [88] were used at all nodes owing to their mitigating effect on hidden state saturation. The design choices were made after conducting ablation experiments which are elaborated in the following section (Ablation). We used a batch size of 300 and a sequence length 150 bins, both of which were observed to be data dependent and the best fit for our data. We used a learning rate of 0.01 for 5 epochs and 0.001 for 5 more epochs. We reinitialized the hidden states of the LSTM after every frame of length 150 and predicted each diagonal block of length 150 with fresh hidden states (Figure 3B). The prediction error improved towards the end of the frame and increased at the start of the next frame (Additional Files: Fig. 1). We tried passing the hidden states across frames and saw that the convergence time significantly increased as the training graph had to be retained across iterations. So we chose to reinitialize the hidden states in each window instead.

We employed PyTorch [89], a Python-based deep learning framework and trained Hi-C-LSTM on GeForce GTX 1080 Ti GPUs with ADAM as the optimizer [90]. All parameters in PyTorch were set to their default values while training. As our primary goal was not to infer values for unseen positions but to form reliable representations for every chromosome, we trained our model on all the chromosomes. For our Hi-C reproduction evaluation, we trained the representations on all chromosomes but the decoders only on a random subset. We chose to train the decoders on a random subset of chromosomes to prevent the decoder from overpowering the representations.

### Hyperparameter selection

To choose the representation size of our model, we performed an ablation analysis. We computed the average mAP across all downstream tasks with the Hi-C-LSTM model which consists of a single layer, unidirectional LSTM with layer norm in the absence of dropout [91] for odd chromosomes and used the even chromosomes to validate whether the choice of hyperparameters remained the same irrespective of chromosome set. We observed the mAP (section Classification) of the Hi-C-LSTM vs. increasing representation size along with Hi-C-LSTM that is bidirectional, in the presence of dropout, without layer norm and 2 layers (Additional Files: Fig. 2). While both the presence of dropout and the absence of layer norm adversely affected mAP, the addition of a layer and a complimentary direction did not yield significant improvements in downstream performance. We conducted a similar ablation experiment and computed the average Hi-C R-squared for the predictions with increasing representation size (Additional Files: Fig. 2) and observed that the performance trend is preserved, which was indicative of the fact that recreating the Hi-C matrix faithfully aids in doing well across downstream tasks. These results were verified to be true for even chromosomes as well (Additional Files: Fig. 2). For both odd and even chromosomes, even though the Hi-C prediction accuracy increased substantially with hidden size, we noticed the elbow at a representation size of 16 for average mAP and therefore set our representation size to that value as a trade-off.

### Hi-C reproduction evaluation

We investigated three hypotheses with following analysis. First, we asked whether the Hi-C-LSTM representations faithfully construct the Hi-C matrix. Second, whether the Hi-C-LSTM representation and the decoder are both powerful in generating the Hi-C map. Third, we evaluated the utility of the representations to infer a replicate map. In all cases, we computed the average prediction accuracy in reconstructing the Hi-C contact matrix, measured using R-squared, which represents the proportion of the variance of the original Hi-C value that’s explained by the Hi-C value predicted by the Hi-C-LSTM.

In our first experiment, we trained both the representations and decoders on replicate 1 (Figure 3A). We took representations trained using all chromosomes from Hi-C-LSTM, SCI and SNIPER and coupled these with some selected decoders, namely, a LSTM, a convolutional neural network (CNN) and a fully connected (FC) feed-forward neural network (used by SNIPER). We compared LSTM with CNN and FC decoders mainly because CNNs provided us with an alternative way of incorporating structure (using moving filters) and FC networks did not include any information about underlying structure. We re-trained these decoders using either of the representations as input, with a small subset of the chromosomes and tested on the rest. All the decoders were configured to have the same number of layers and hidden size per layer. As the decoders were separately trained, this process allowed us to check the power of the representations alone, moreover, as a small subset of chromosomes were used to train the decoder, we reduced the possibility of the decoders overfitting.

In our second experiment (Figure 3B), we trained the representations on replicate 1 using all chromosomes, and repeated the aforementioned decoder training process on replicate 2.

### Comparison methods

We compared our downstream classification results with five alternatives: two variations of SNIPER [92], one with inter-chromosomal (SNIPER-INTER) and the other with intra-chromosomal contacts (SNIPER-INTRA), SCI [93] and two baselines, namely, the subcompartment-ID (SBCID) and principal component analysis (PCA). SNIPER-INTRA was the same as the original SNIPER-INTER, modified to take the intra-chromosomal row as input instead of the inter-chromosomal row. All the parameters for the two SNIPER versions and SCI were set as given in their respective papers [11], [12]. The SBCID baseline used the one-hot-encoded vector of the subcompartment as the representation at the position under contention. The PCA baseline assigned the principal components from the PCA of the Hi-C matrix as the representations.

### Element identification evaluation

We used the following analysis to evaluate the ability of a representation to identify genomic phenomena and chromatin regions.

For each type of element, a boosted decision tree classifier called XGBoost [41] was trained on the representations. We employed tree boosting as it is shown to outperform other classification models with respect to accuracy when ample data is available. Following Avocado [70], we used XGBoost with a maximum depth of 6 and a maximum of 5000 estimators and these parameters were chosen following ablation experiments with odd chromosomes as the training set and even chromosomes as the test set (Additional Files: Fig. 3). N-fold cross-validation, with *n* = 20, was used to validate our training with and an early stopping criterion of 20 epochs. The rest of the XGBoost parameters were set to their default values.

For each task, the genomic loci under contention were assigned labels. All tasks were treated as binary classification tasks, except the subcompartments task, which was treated as a multi-class classification task. For tasks without preassigned negative labels, negative labels were created by randomly sampling genome-wide, excluding the regions with positive labels.

The XGBoost classifier was given the representations at these genomic loci as input and the assigned labels as targets. The classifier was evaluated using the metric of mean average precision (mAP), which is a standard metric for classification tasks and is defined as the average of the maximum precision scores achieved at varying recall levels.

### Sequence attribution

We validated the utility of the Hi-C-LSTM representations in locating genomic regions important for conformation using feature attribution analysis. Feature attribution was carried out on the intra-chromosomal representations using Integrated Gradients [94]. Integrated Gradients is a feature attribution technique that follows an axiomatic approach to attribution, adhering to the axioms of sensitivity and implementation invariance. Sensitivity implies that if the input and baseline differs in one feature and have different predictions, then the differing feature should be assigned a non-zero attribution. Implementation invariance requires that two networks, whose output is equal for every input despite having different implementations, should have the same attributions. We used Captum [95], a Integrated Gradients feature attribution framework in PyTorch that is generic and works with sequential models. The resulting feature attributions were summed across all features, giving us one importance score for every position in the genome. The feature importance scores were then subjected to log normalization followed by min-max normalization (Eq. 5). Specifically, let *IG* be to the integrated gradients (IG) score, *IG*_*min*_ and *IG*_*max*_ be the minimum and maximum IG scores. The normalized IG score *IG*_*norm*_ is defined as

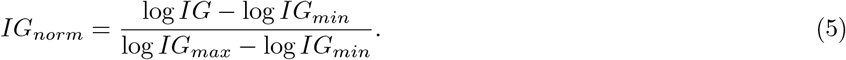

### In-silico perturbation

The Hi-C-LSTM enables us to perform in-silico deletion, orientation replacement and reversal of genomic loci and predict changes in the resulting Hi-C contact map. We performed three types of experiments:: knockout, CTCF orientation replacement, and duplication. In a knockout experiment, we chose certain genomic sites (such as CTCF and cohesin binding sites) and replaced their representations with a null representation. As a null representation, we used the average representations in a window of 0.2 Mbp around the site in question, because this captures the genomic neighborhood while removing the features specific to site. The knockout of the representation at a particular row affects not just the Hi-C inference at columns corresponding to that row but also the succeeding rows because of Hi-C-LSTM’s sequential behavior. The LSTM weights remain unchanged, but as the input to the LSTM is modified, the inferred Hi-C contact probability is altered based on the information retained by the LSTM about the relationship between the sequence elements under contention and chromatin structure.

In a CTCF orientation replacement experiment, we replaced the representations of downstream-facing CTCF motifs with the genome-wide average of the upstream-facing motifs and vice versa. This was done under the assumption that the average representation of the given orientation would encapsulate the important information regarding the role played by the orientation in chromatin conformation.

Our duplication experiment was carried out by creating a tandem duplication the representations from the 2.1 Mbp region between 67.95 Mbp to 70.08 Mbp in chromosome 7 region [56] and then passing the resulting representation matrix to the LSTM to infer contacts. Given our Hi-C resolution of 10 kbp, the duplicated region corresponds 214 bins, i.e., bin 6795 to bin 7008. Specifically, the duplicated representation matrix 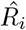 is defined as 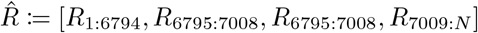.

To enable comparison to Hi-C data mapped to the original pre-duplication reference genome, we combined inferred contacts from both copies. This combination is required because Hi-C reads cannot be disambiguated between the two duplicated copies when they are mapped to the reference genome. Specifically, we passed the predicted contact probability *cp* through the inverse exponential transformation to define predicted read counts 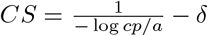 (see Eq. 1). We summed predicted read counts from the two duplicated copies to simulate mapping reads from both copies to the same reference genome *CS′*, then re-applied the exponential transform to obtain predicted contact probability *cp′*.

Our baseline for the quantitative evaluation was the original pre-duplication Hi-C for the interactions between the upstream, downstream and duplicated regions, and the genomic average for the interactions of the duplicated region with itself. We considered a window of 214 bins (length of the duplicated region), and computed the average genomic contact strength for the bins with themselves in a window of this size.

## Declarations

### Availability of data and materials

The data that support the findings of this study are publicly available to download and are referenced in the bibliography. Refer to Methods for more details. The data and representations generated from the project can be found at [60].

### Competing interests

The authors declare that they have no competing interests.

### Funding

This work was funded by NSERC Discovery Grant awards RGPIN-2015-03948 and AWD-001606, a Simon Fraser University President’s Research Startup Grant, and a Four Year Doctoral Fellowship from the University of British Columbia.

### Authors’ contributions

K.B.D. led ideation, genomic data processing, building and validating the machine learning model, wrote the first draft of the manuscript, and edited the manuscript. A.M. contributed towards ideation, data processing, parallelization of the model and model validation. V.K.B. partially funded the project. M.W.L. supervised the project. All authors participated in the design of the study, the interpretation of results, and editing the manuscript. All authors read and approved the final manuscript.

### Ethics approval and consent to participate

Not applicable.

### Consent for publication

Not applicable.

### Authors’ information

K.B.D. is a PhD student at UBC where he works on computational genomics jointly under the information theory group at UBC and computational biology group at The Simon Fraser University. His work focuses on building machine learning tools to aid in the understanding of structural and functional genomic data. A.M. holds an MSc in Bioinformatics from the University of British Columbia and is currently a PhD student at Simon Fraser University where she works in the computation biology group. Her research focuses on the use of machine learning approaches in the analysis for genomics data. E.A-J. is a PhD student at Imperial College London studying 3D genome organisation and gene expression. M.M. is a Career Scientist at the MRC’s Clinical Sciences Centre at Imperial College. He is a molecular immunologist whose work has been central to the understanding of development, and cellular reprogramming. V.K.B. is a Fellow of the IEEE, The Royal Society of Canada, and currently a Professor in the Department of Electrical and Computer Engineering at the University of British Columbia in Vancouver. M.W.L. is an Assistant Professor in Computing Science at Simon Fraser University where his research focuses on developing machine learning methods applied to high-throughput genomics data sets.

## Additional Files

**Additional file 1** — Supplementary Results

The additional file contains supplementary figures of salient features of Hi-C-LSTM predictions, ablation experiments with Hi-C-LSTM, parameter search for the XGBoost classifier, confusion matrix for classification of subcompartments, and feature importance for Segway-GBR labels.

## Notes

https://github.com/smaslova/HiCLSTM

### Competing Interest Statement

The authors have declared no competing interest.

