## Supplementary Material for "Hi-C-LSTM: Learning representations of chromatin contacts using a recurrent neural network identifies genomic drivers of conformation"

### Additional File 1

This file contains 5 supplementary figures pertaining to ablation experiments, model parameter choices and model performance.

---

- Fig. 1 points out the salient features of Hi-C-LSTM predictions.
- Fig. 2 shows the ablation experiments carried out to determine the representation size of Hi-C-LSTM.
- Fig. 3 shows the results from the parameter search for the XGBoost classifier.
- Fig. 4 displays the confusion matrix for the subcompartment classification task.
- Fig. 5 shows the feature importance scores for domain labels given by Segway-GBR.

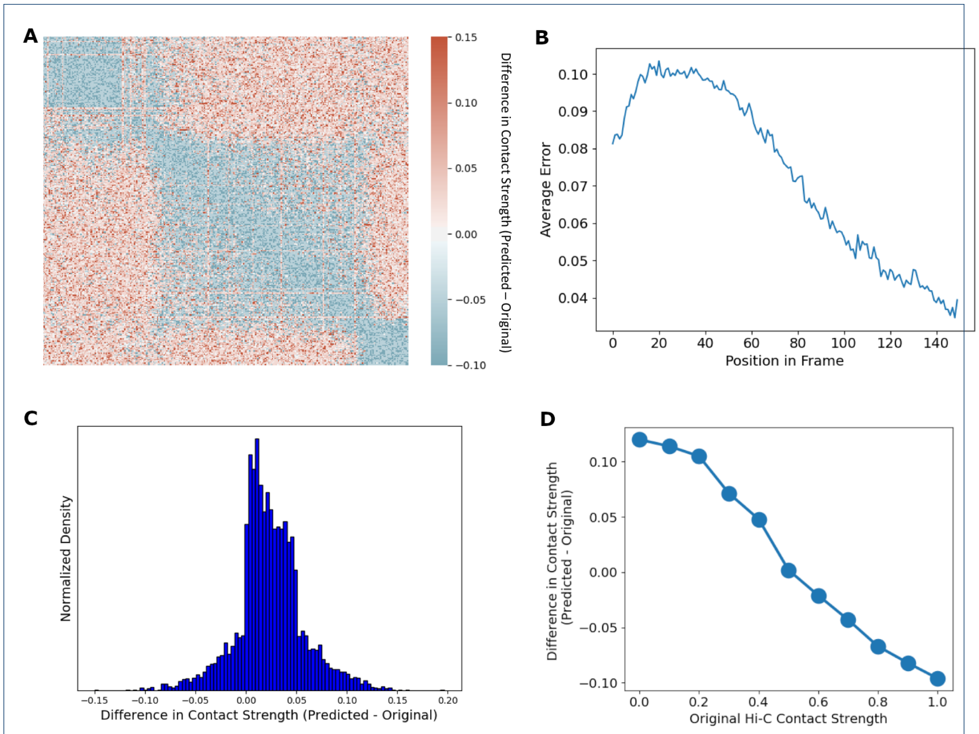


Figure 1: Salient characteristics of the Hi-C-LSTM predictions are analysed by comparing them with original Hi-C values and visualizing their range. **A**) The difference in contact strength between the predicted Hi-C map and original Hi-C map for a selected genomic region between 41.4 to 41.6 Mbp. **B**) Average prediction error within each frame. We see that error is high at the beginning of the frame and improves towards the end of the frame, explaining the discontinuity in our predictions at the start of each frame. **C**) The histogram of the difference between predicted and original Hi-C values. The y-axis shows the normalized density and the x-axis refers to the difference in contact strength between predicted and original Hi-C. The plot reveals the low spread and slight positive skew of prediction errors. **D**) The variation of difference in predicted and original Hi-C (y-axis) across the range of original Hi-C values (x-axis). Its seen that the errors go from positive to negative as values of original Hi-C increase, which can also be seen in **A**, pointing to the Hi-C-LSTM over-predicting low values and under-predicting high values of Hi-C.

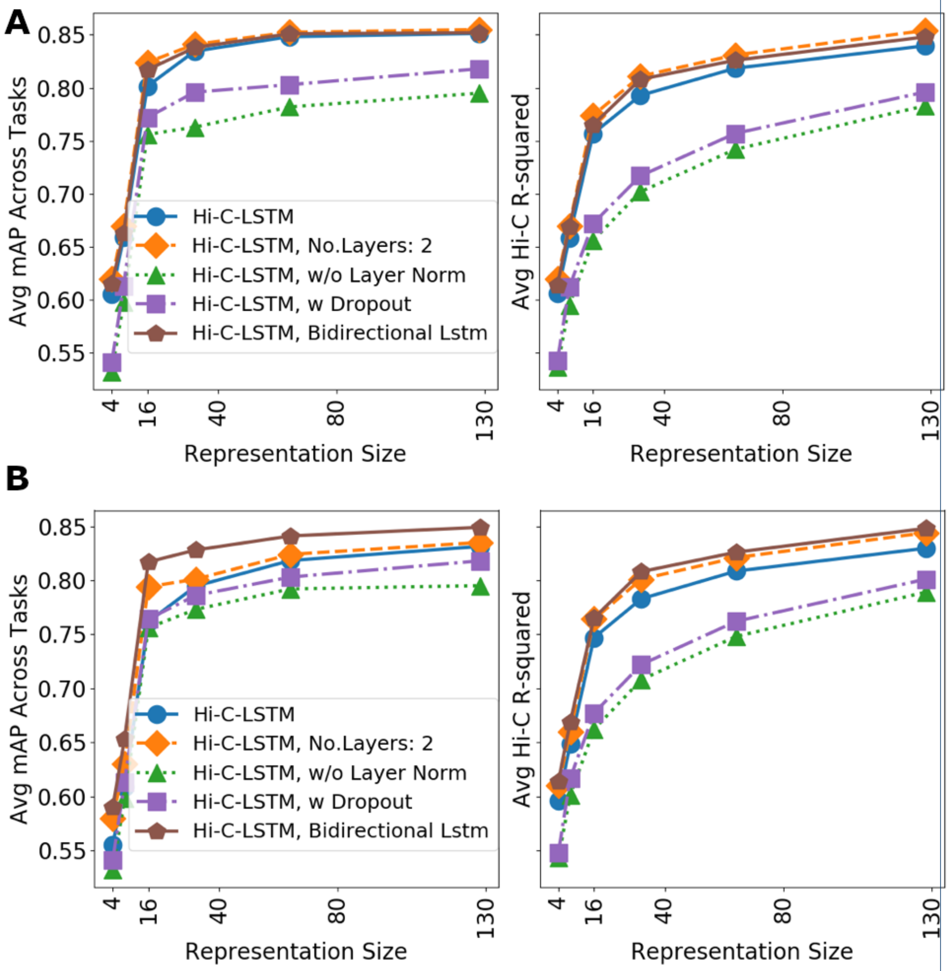


Figure 2: **Ablation Results**: To choose the representation size of our model, we performed an ablation analysis. **A:L**) Average mAP across downstream tasks (y-axis) with the Hi-C-LSTM model (single layer, unidirectional LSTM with layer norm in the absence of dropout) and its ablations with increasing representation size (x-axis) for odd chromosomes. **A:R**) The average Hi-C R-squared (y-axis) with representation size (x-axis) for odd chromosomes. The ablations of the model are shown in the legend with different markers and colors. The performance trend is preserved across **A:L** and **A:R**, which is indicative of the fact that faithfully creating the Hi-C matrix helps in performing well across classification tasks. **B**) Similar trend is seen for even chromosomes. In both **A** and **B**, the Hi-C R-squared increases considerably with hidden size, however, we notice an elbow at an embedding size of 16 for average classification mAP and thus set our embedding size to that value as a trade-off. It is interesting to see that a low-dimensional Hi-C-LSTM representation is able to successfully create the H-C matrix as well as do well across classification tasks.

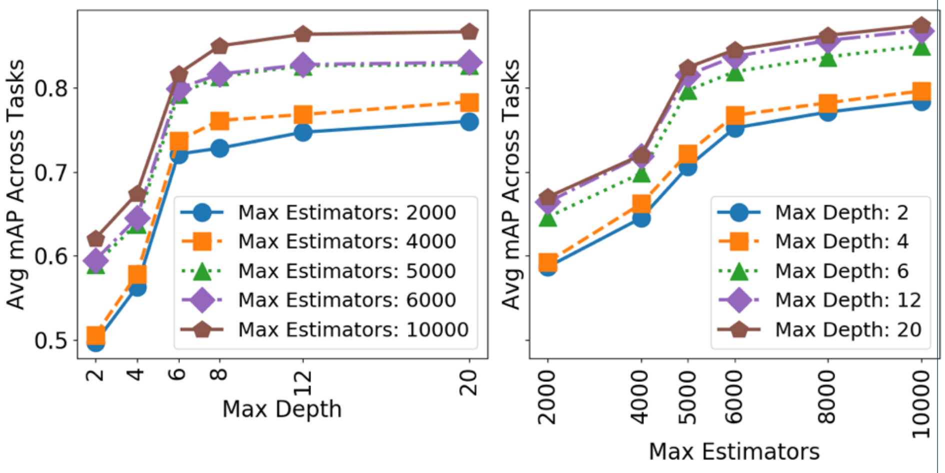


Figure 3: **XGBoost Parameters**: To choose the maximum depth and the maximum number of estimators of the XGBoost classification framework, we performed an ablation analysis with odd chromosomes as the training set and even chromosomes as the test set. The test results are shown here. **A**) Average mAP across downstream tasks (y-axis) using the Hi-C-LSTM representations with increasing maximum depth (x-axis). The legend shows the maximum number of estimators used in each case. As the elbow in **A** is seen at a maximum depth of 6 for all values of maximum estimators, we set our maximum depth to this value, achieving optimum gains in performance and run time. **B**) The average mAP across downstream tasks (y-axis) with increasing maximum number of estimators (x-axis). The legend shows the maximum depth used in each case. In **B**, the elbow is clearly seen at 5000 maximum estimators for different maximum depths. Following this, we set our maximum number of estimators to 5000.

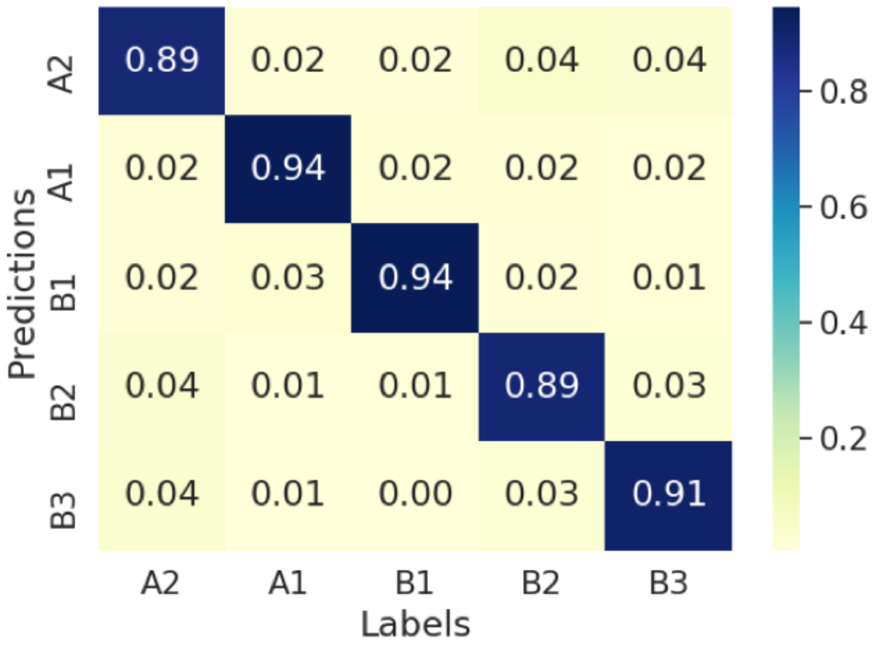


Figure 4: **Subcompartment Confusion Matrix**: The Hi-C-LSTM representations are used to classify subcompartments in the Genome and the resulting confusion matrix is plotted. The x-axis shows the subcompartment labels. The y-axis shows the subcompartment predictions by the XGBoost classifier, and the values in the confusion matrix are true prediction percentages. All subcompartments achieve a true prediction percentage $>\approx 90\%$. The A1 and B1 subcompartments achieve the highest true predictions while the other subcompartments follow closely.

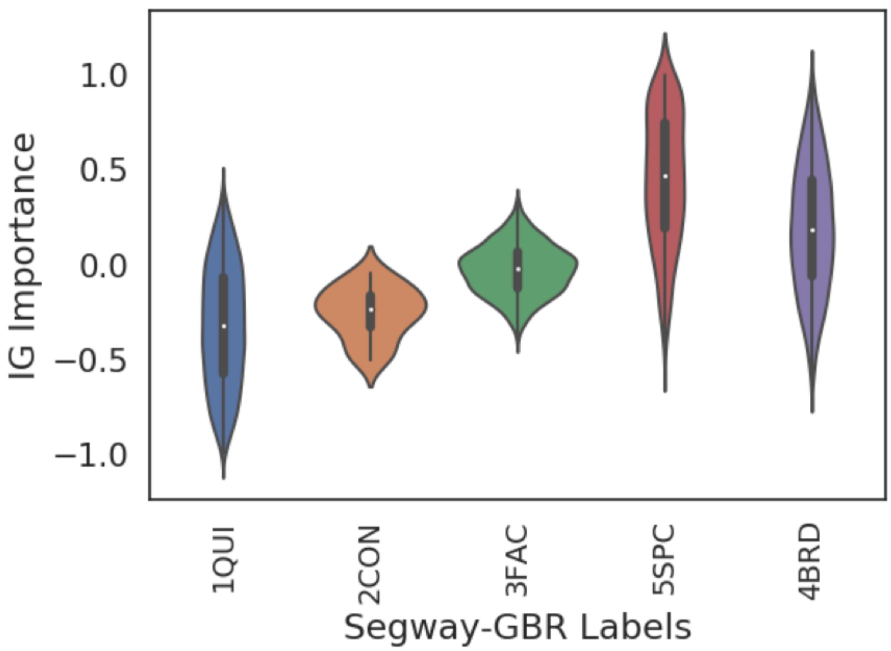


Figure 5: **Segway-GBR Feature Importance**: Segway, by itself, cannot handle chromatin conformation data, however, when coupled with graph-based regularization (GBR), a method that encourages positions that are close in 3D space to occupy the same type of domain by including pairwise prior information during genome annotation, a model revealing known chromatin domains is obtained. Segway GBR gives five types of domains: quiescent (QUI), constitutive heterochromatin (CON), facultative heterochromatin (FAC), broad expression (BRD), and specific expression (SPC). Quiescent, constitutive and facultative domains are repressive. Broad and specific expression domains are active. Unlike BRD domains, genes present in a SPC domain are highly expressed compared to their mean in all cell types, hinting that SPC domains might be highly activating. The plot of aggregated feature importance scores shows largely positive values for SPC and BRD domains and largely negative values for QUI and CON domains, while FAC domains exhibit a bit of both. Both the highly activating nature of SPC domains and the repressive nature of QUI and CON domains is thereby validated.
